# Mitigating biomass composition uncertainties in flux balance analysis using ensemble representations

**DOI:** 10.1101/652040

**Authors:** Yoon-Mi Choi, Dong-Hyuk Choi, Yi Qing Lee, Lokanand Koduru, Nathan E. Lewis, Meiyappan Lakshmanan, Dong-Yup Lee

## Abstract

The biomass equation is a critical component in genome-scale metabolic models (GEMs): it is used as the *de facto* objective function in flux balance analysis (FBA). This equation accounts for the quantities of all known biomass precursors that are required for cell growth based on the macromolecular and monomer compositions measured at certain conditions. However, it is often reported that the macromolecular composition of cells could change across different environmental conditions; the use of the same single biomass equation in FBA, under multiple conditions, is questionable. Thus, we first investigated the qualitative and quantitative variations of macromolecular compositions of three representative host organisms, *Escherichia coli*, *Saccharomyces cerevisiae* and *Cricetulus griseus*, across different environmental/genetic variations. While macromolecular building blocks such as DNA, RNA, protein, and lipid composition vary notably, variations in fundamental biomass monomer units such as nucleotides and amino acids are not appreciable. We further observed that while macromolecular compositions are similar across taxonomically closer species, certain monomers, especially fatty acids, vary substantially. Based on the analysis results, we subsequently propose a new extension to FBA, named “Flux Balance Analysis with Ensemble Biomass (FBAwEB)”, to embrace the natural variation in selected components of the biomass equation. The current study clearly highlights that certain components of the biomass equation are very sensitive to different conditions, and the ensemble representation of biomass equation in the FBA framework enables us to account for such natural variations accurately during GEM-guided *in silico* simulations.

## 1. Introduction

Constraint-based metabolic reconstruction combined with flux balance analysis (FBA) is a popular approach for analysing cellular metabolic behaviours *in silico* (Bordbar *et al*., 2014). Unlike dynamic modelling which requires detailed kinetic parameters, it simply uses information on metabolic reaction stoichiometry and mass balances around the metabolites, under pseudo-steady state assumption (Lewis *et al*., 2012). Such simplicity of FBA and massive genomic data available from public databases e.g., NCBI Assembly database (Kitts *et al*., 2016), enabled the development and use of hundreds of genome-scale metabolic models (GEMs) for a multitude of species across all three domains of life (Yasemi and Jolicoeur, 2021). These GEMs have been successfully applied in various studies including microbial evolution, metabolic engineering, drug targeting, context-specific analysis of high throughput omics data and the investigation of metabolic interactions among cells and/or organisms (Gu *et al*., 2019).

Basically, FBA is an optimization-based approach where a particular cellular objective is maximized or minimized while simultaneously constraining the mass balance, thermodynamic and enzyme capacity of a metabolic network to determine the plausible steady-state fluxes. Of several objective functions that have been considered to interrogate the metabolic states and cellular behaviours, the maximization of biomass production has been most commonly adopted in FBA with a principal hypothesis that living cells typically strive to grow as fast as possible, at least under their exponential growth phase (Feist and Palsson, 2010; Schuetz *et al*., 2007). Therefore, all reconstructed GEMs provide an artificial reaction, referred to as the ‘biomass equation’, that accounts for the stoichiometric proportions of various compounds that make up macromolecules of the cellular biomass, e.g., protein, DNA, RNA, carbohydrate, and lipid. It should be noted that the macromolecular and monomer compositions in the biomass equation are empirically determined under a certain experimental condition and assumed to remain similar across a wide range of growth environments. In addition, wherever necessary, biomass equations also often use data obtained from taxonomically close organisms since it is assumed that the macromolecular compositions remain similar across closely related species. However, it is well-documented that cellular volume and the compositions of macromolecular components may vary depending on growth conditions and/or genetic makeup of cells (Volkmer and Heinemann, 2011; Beck *et al*., 2018; Széliová *et al*., 2020; Schinn *et al*., 2021). For example, the RNA/protein ratio of *Escherichia coli* exhibits robust correlation with their growth phase and culture conditions such as nutrient utilization/depletion and waste-product accumulation (Scott *et al*., 2010). Recently, other studies also demonstrated the cell line-dependent variation in protein and lipid composition in immortalized mammalian cell lines derived from the same original tissue (Széliová *et al*., 2020) and large variations in cell size across species (Westoby *et al*., 2021).

Since almost all FBA simulations involve the growth maximization, the perseverative use of a biomass equation formulated from a single compositional dataset raises such arguable question about its impacts on the model predictions in the research community. In this regard, previous studies have examined effects of variations in individual biomass components on phenotype predictions (Feist *et al*., 2007; Dikicioglu *et al*., 2015; Koduru *et al*., 2017; Xavier *et al*., 2017; Schinn *et al*., 2021). While these studies showed the variable nature of biomass and its impact on FBA results, a consensus on the use of the same biomass equation across multiple conditions as well as the data source used to draft such reactions is yet to be reached: Do all macromolecular and monomer compositions in biomass vary across environmental conditions? If so, how significant are those variations? Do phylogenetically close organisms have similar macromolecular compositions? How reliable is the estimation of biomass composition from omics datasets? How much does such a natural variation of biomass composition impact model predictions?

To address the above-mentioned questions, here, we first examined the variations in biomass compositions in three representative host organisms, namely *E. coli*, *Saccharomyces cerevisiae* and Chinese hamster ovary (CHO) cells. We also investigated the macromolecular heterogeneity between these species and their phylogenetically close organisms and analysed the quantitative variation between monomer compositions obtained from omics datasets and the experimentally measured ones. Based on the analysis results, we newly incorporated an ensemble of biomass equations within the FBA framework, to better capture the natural variation in biomass compositions.

## 2. Results

### 2.1 Assessing the validity of various common assumptions made while drafting the biomass equation

Typically, drafting a biomass equation in GEMs begins with the measurement of weight fraction of macromolecular components (protein, DNA, RNA, carbohydrate, lipid, and cofactors), which together make up 1 gram of dried cells. Subsequently, the quantity of monomer metabolites which make up each of these macromolecular classes are also measured. For example, the proportion of different amino acids which constitute one gram, or one mole of total protein is needed to draft the biomass equation. Here, it should be emphasized that most published GEMs derive this information (macromolecular and monomer quantifications) from a single experimental condition, typically under exponential growth phase. As such, one key requirement for the validity of the biomass equation, which is drafted based on a single experimental condition, is that the macromolecular and monomer components of the cells should remain reasonably robust across different conditions. However, contrary to such an assumption, it has been often reported that the cellular compositions and energy requirements change across different environmental conditions even in the same organism (Scott *et al*., 2010; Széliová *et al*., 2020; Lahtvee *et al*., 2016). Therefore, understanding the natural variability in the cellular compositions is *a priori* while drafting biomass equations.

We hereby analysed the variations in macromolecular and monomer compositions of three highly divergent and representative organisms, *E. coli* (a prokaryote), *S. cerevisiae* (a unicellular eukaryote), and CHO cells (immortalised cells derived from a multicellular eukaryote) compiled from cells cultured under various environmental and/or genetic backgrounds (see **Material and Methods**). To compare the variability of individual biomass components, we calculated the coefficient of variation (CV), which is defined as a ratio between the sample standard deviation and the sample mean, or a percentage (%). Such comparison revealed that macromolecules show larger variation than monomers (CV = 6-87% compared to that of 2-50%). Among various macromolecules, while protein had the smallest CV% in all three organisms, other macromolecules surpassed 20% of CVs in most cases (**Figure 1a**). Notably, lipids, biomolecular compounds which are involved in long term energy storage and cellular membrane reconstitution, have substantially higher variability than other macromolecules in general. We also observed that all macromolecules showed increasing CVs in sequence of *E. coli*, *S. cerevisiae*, and CHO cells (**Figure 1a**). Since prokaryotic cells are more adept in adapting to various niches through their faster exchange of various molecules with adjacent environments owing to a small cellular size with high surface area, their metabolism as well as cell growth can be stimulated much faster than eukaryotes (Sonea and Mathieu, 2000). This could result in a faster turnover of cellular components, e.g., macromolecular composition (Sonea and Mathieu, 2000; Finkel *et al*., 2016). Among monomers, while amino acids and nucleotides showed relatively less variability, fatty acids showed very large variations across different growth conditions (**Figure 1b**). The high variability of fatty acids distributions is generally attributed to their adaptations against perturbed environments, e.g., temperature, pH, salt, or dietary conditions (Guerzoni *et al*., 2001; Prakash *et al*., 2015; Levental *et al*., 2020). Overall, the comparisons of biomass compositions clearly showed that while monomer compositions except fatty acids are relatively stable, compositions of macromolecules vary considerably depending on the genetic and environmental background.

**Figure 1.**
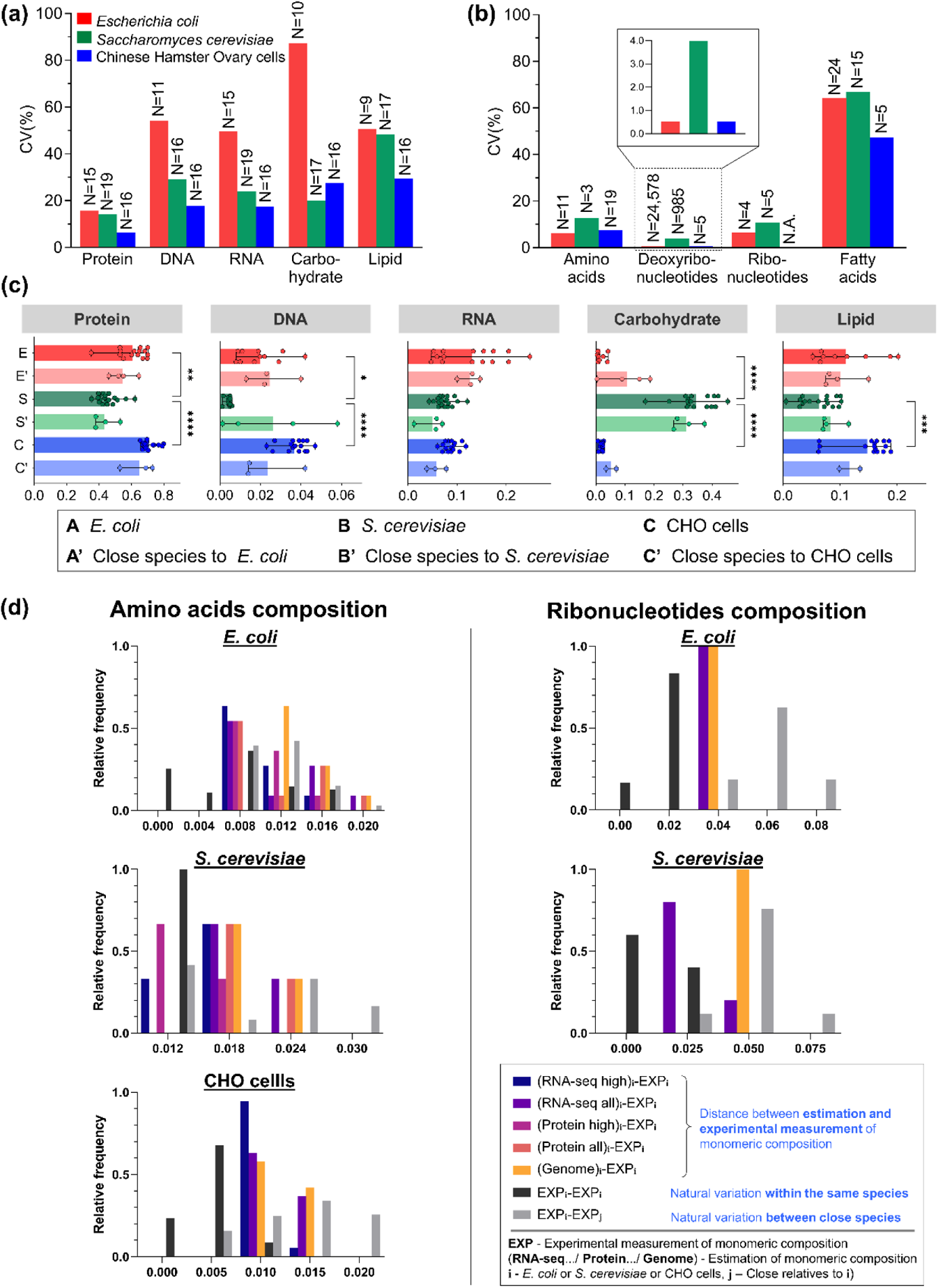
Inherent variations in macromolecules within/across species and monomers within the same species. **1a,b**. The coefficient of variations (CVs) of five macromolecules classes within the same species (**a**) and 4 monomer groups (**b**) within each of *E. coli*, *S. cerevisiae*, and CHO cells. **1c**. Macromolecular compositions across phylogenetically close species including Gram-negative bacteria, yeast species, and murine cells or *M. musculus.* **1d**. Distance (root mean square deviation, RMSD) between various sources of composition data was compared for amino acid (left) and ribonucleotide (right) composition of *E. coli*, *S. cerevisiae*, and CHO cells. Distance data is not available for ribonucleotides composition of CHO cells.

We next tested the validity of borrowing individual components from phylogenetically closer species to draft the biomass equation. For example, we examined the biomass equations of eight phylogenetically different yeast species: *Candida glabrata* (Xu *et al*., 2013), *Candida tropicalis* (Mishra *et al*., 2016), *Kluyveromyces lactis* (Dias *et al*., 2014), *Pichia pastoris* (Chung *et al*., 2010), *S. cerevisiae* (Mo *et al*., 2009), *Schizosaccharomyces pombe* (Sohn *et al*., 2012), *Scheffersomyces stipitis* (Balagurunathan *et al*., 2012) and *Yarrowia lipolytica* (Kavšcek *et al*., 2015); we found that several components were borrowed from one model to another with *S. cerevisiae* being the common source. Likewise, many other published GEMs have also adopted biomass data from a close species to formulate a biomass equation due to lack of relevant data (Nogales *et al*., 2008; Zhang *et al*., 2009; Aggarwal *et al*., 2011; Ulas *et al*., 2012; Xu *et al*., 2013; Mishra *et al*., 2016; Dias *et al*., 2014; Chung *et al*., 2010; Sohn *et al*., 2012; Kavšcek *et al*., 2015; Koduru *et al*., 2020). Hence, we collected and compared the experimentally measured macromolecular and monomer compositions of phylogenetically close species to all the three species considered here (**Figure 1c, Supplementary file 1**). A wide range of species were included in such comparison instead of constraining into the same genus; several Gram-negative bacteria and yeast species were compared with *E. coli* and *S. cerevisiae*, respectively. In the case of CHO cells, we compared it only with mouse cells due to scarcity of such data from other higher eukaryotes. Each of the three organisms considered here have similar macromolecular distributions with their phylogenetically closer organisms. At the same time, we also noted significant macromolecular composition differences between various classes of species; biomass of Gram-negative bacteria was vastly different from yeasts and mammalian cells. It should be highlighted that while most macromolecular compositions were similar to their phylogenetically closer organisms, substantial differences were observed in the carbohydrate composition between *E. coli* and its close species. As this macromolecule was highly variable even within the same species under various environmental conditions (**Figure 1a**), care should be taken when borrowing carbohydrate data (mass%) from close species. We further compared the amino acids and fatty acids across species to evaluate the differences or similarities in monomer compositions. We were unable to compare nucleotide composition across species due to lack of such experimental data. The composition of both amino acids and fatty acids was markedly different across species unlike macromolecules (**Supplementary file 1**). Overall, we conclude that although the macromolecular contents vary across environmental conditions to a certain extent, the relative composition of each component is similar to the phylogenetically closer species and can be borrowed when the relevant data is not available. However, this assumption is not true for monomers; amino acid and fatty acid data cannot be borrowed from phylogenetically/taxonomically closer species and need to be experimentally determined.

It has been previously suggested that amino acid and nucleotide composition can be estimated from genome (Thiele and Palsson, 2010) or transcriptome (Santos and Rocha, 2016) datasets. Hence, we next evaluated whether the “omics” data can be used while drafting biomass equations. We first estimated the amino acid composition from transcriptome (mRNA-seq or rRNA depleted) and proteome datasets, and ribonucleotide composition from whole transcriptome datasets of *E. coli*, *S. cerevisiae* and CHO cells (see **Material and Methods**). We then evaluated the “closeness” between the experimentally measured amino acid and ribonucleotide composition and the “-omics” estimated ones using the Euclidean distance – a measures of divergence between two datasets (see **Material and Methods**). In regards to ribonucleotides, we calculated the distance between omics (genome and transcriptome) data derived compositions and experimentally measured ones (genome_n_ – experimental_n_ and transcriptome_n_ – experimental_n_) and compared it with the distance between experimentally measured composition from the same organism and its closer relatives (experimental_n_ – experimental_m_). This comparison revealed that while no significant difference was observed between genome-derived estimates and measurements from closer organisms, the transcriptome-derived estimates had smaller distances than the closer organisms, indicating that transcriptome can be reliably used to estimate ribonucleotide distribution rather than borrowing it from closer organisms. Next, we calculated the amino acid composition from omics (genome, transcriptome and proteome) data and compared it with experimentally measured ones. Note that we used two combinations of transcriptome and proteome data, all and highly expressed, since it has been previously suggested that the highly expressed (top 10%) mRNA transcripts and proteins account for large proportion of the overall transcriptome and proteome outputs, respectively (genome_n_ – experimental_n_, transcriptome-all_n_ – experimental_n_, transcriptome-high_n_ – experimental_n_, proteome-all_n_ – experimental_n_ and proteome-high_n_ – experimental_n_). We then compared these Euclidean distances with the one calculated to estimate the divergence between the organism and its relatives (experimental_n_ – experimental_m_). The distances between amino acid composition estimated from highly expressed transcripts/proteins were significantly less than that of experimental measurements from closer organisms (**Supplementary table 1**), and thus highlighting the estimation of amino acid composition from highly expressed transcripts/proteins is a good choice.

### 2.2 Sensitivity of macromolecular/monomer composition in phenotype predictions

We next examined the sensitivity of predicted *in silico* growth rates and intracellular metabolic fluxes upon varying the macromolecular and monomer compositions in all three species across diverse environments, which allowed us to identify the key biomass components with the greatest influence on FBA results. Particularly, the sensitivity analysis was carried out under aerobic and anaerobic conditions in *E. coli*, in three different carbon sources (glucose, xylose and ethanol) in *S. cerevisiae* and three different cell lines in CHO cells. We used the recently published GEMs of each organism *i*ML1515 (Monk *et al*., 2017), Yeast8.0.0 (Lu *et al*., 2019), and *i*CHO2291 (Yeo *et al*., 2020) to analyze the flux sensitivities. Since the *E. coli ‘*core’ biomass equation was used as a template which does not have carbohydrate content (Monk *et al*., 2017), the sensitivity of carbohydrate in *E. coli* was excluded in this analysis.

Initially, we examined the sensitivity of macromolecular and monomer composition on growth rate predictions. Our analysis indicated that protein and lipid compositions were the most sensitive among different macromolecules while DNA was found to be the least sensitive (**Supplementary Figure 1**). Such a trend is expected since proteins and lipids have the largest fraction (by mass) in dry cell weight while the composition of DNA is almost negligible. Our analysis also unravelled condition specific sensitivities of macromolecular composition in FBA. For example, the sensitivity of macromolecules predicted under aerobic condition was higher than that of anaerobic conditions in *E. coli*. Similarly, the sensitivities were lower in reduced substrate ethanol than that of glucose and xylose. Among monomers, only amino acid composition was observed to be significantly sensitive in growth rate predictions.

Subsequently, we explored the effect of macromolecular and monomer compositions on internal flux distribution of all the reactions using pFBA. The flux sensitivity was quantified based on Euclidean distance between the predicted fluxes in the reference state, i.e., original biomass equation, and those in a perturbed state, with the only difference between the two states being the biomass composition (see **Material and Methods**). Similar to the variations in growth rates, intracellular fluxes were highly sensitive to protein and lipid macromolecule compositions in all three model organisms (**Supplementary Figure 2**). With regards to monomer, we observed that the resultant flux to be highly dependent on amino acid compositions, while fatty acid composition affects flux distributions in CHO cells but not in others. Such variations in the intracellular fluxes across all reactions clearly indicate that the biomass composition is most critical for predicting accurate internal flux distributions. While we have assessed the sensitivity of biomass components on internal flux distributions using pFBA, this observation should hold true even for other constraint-based analysis such as flux variability analysis (Mahadevan and Schilling, 2003) and Monte Carlo flux sampling (Schellenberger and Palsson, 2009).

Here, it should be highlighted that since the absolute sensitivities across the three models had different levels of magnitude, we normalized the sensitivity values with respect to the sum of the absolute sensitivities of biomass components to understand the relative influence of each component on flux prediction. We also performed a statistical evaluation for the sensitivities of macromolecules and monomers to determine the significance differences between them (see **Material and Methods**). In general, macromolecules showed significantly higher sensitivities than monomers (q<0.05; adjusted p values) (**Supplementary Figure 3**). Among different macromolecules, protein and carbohydrate showed larger normalised sensitivity than any others (**Figure 2**). The CHO model showed very high absolute (flux) sensitivity of protein over two other models, which was consistent with a large difference in the predicted growth rate (**Supplementary figure 1**). Another common result observed across the three GEMs was that the flux sensitivities upon changing DNA composition were almost negligible. The flux sensitivities of monomers including fatty acids were mostly insignificant compared to macromolecules with exception of CHO cells.

**Figure 2.**
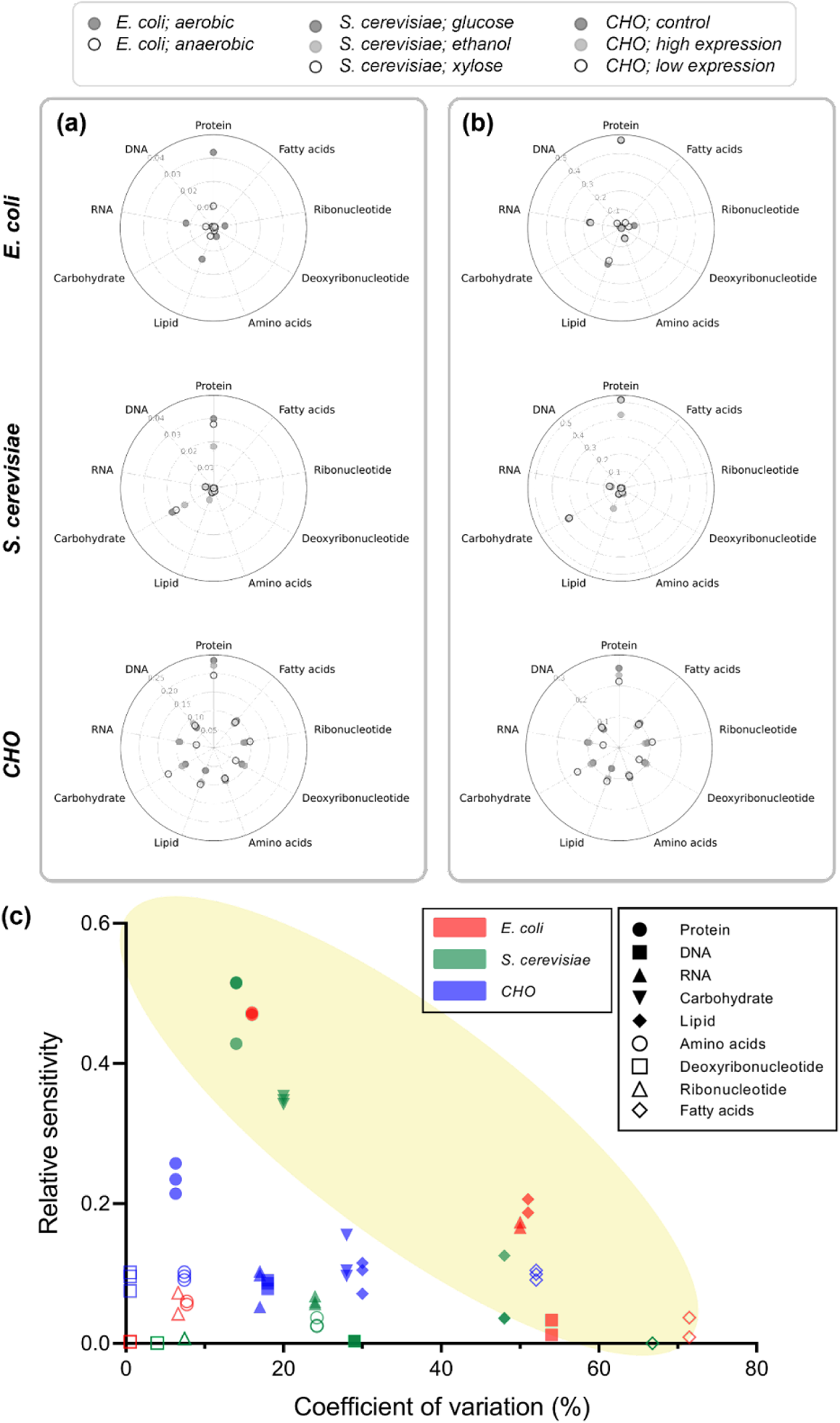
Sensitivity and variability of biomass components. **2a,b**. Absolute (**a**) and relative (**b**) sensitivities of biomass components evaluated under various conditions. A relative sensitivity was obtained by normalizing an absolute sensitivity by the sum of the absolute sensitivities of all macromolecules and monomers for every model condition. **2c**. Distribution of biomass components along the coefficient of variation and relative sensitivity determined under various conditions.

Finally, by aggregating our results, we propose to label a certain biomass component “critical” in terms of its potential impact on model predictions, if it falls in any of the following three categories: 1) high CV & high sensitivity 2) high CV & low sensitivity 3) low CV & high sensitivity. Briefly, we consider a critical biomass component for flux predictions when the component has either high CV or high sensitivity or both. For example, a biomass component with high natural variability but low sensitivity, e.g., fatty acids, can result in more significant changes in flux prediction because we only varied the components to a 25% limit in the sensitivity analysis. On the other hand, components with small CV and high sensitivity, e.g., protein, should also be considered as critical since they may lead to large differences in flux predictions even with small measurement errors (**Figure 2c**). Overall, we observed that accurately accounting for macromolecules such as protein, RNA, carbohydrate, and lipid, and the fatty acid monomer is significant to formulate the biomass equation.

### 2.3 Accounting for biomass natural variation by “Flux Balance Analysis with Ensemble Biomass (FBAwEB)”

As we have shown in the previous sections, while some biomass components naturally vary across different conditions, some others are highly sensitive to FBA results, and thus the use of a biomass equation drafted from a single experimental condition could be inaccurate. To address both the natural variation in macromolecular components and potential experimental errors in the measurement of sensitive components, we propose a new method called “Flux Balance Analysis with Ensemble Biomass (FBAwEB)” which represents biomass equation as ensembles within the FBA framework (**Figure 3**). In this approach, the biomass components that naturally vary or are highly sensitive to FBA results are sampled “n” number of times within the known range and “n” biomass reactions with various combinations of the diversified biomass composition were generated and normalized to make up 1 gram of cell in total. Here, “n” is an arbitrary number, and we have chosen 5000 to account for a good mix of various biomass components within the varying range. Subsequently, FBA/pFBA is implemented “n” times with each of the newly generated “n” biomass equations as the objective function. The distribution of fluxes obtained from “n” number of simulations are then analysed to extract the plausible flux ranges for each reaction, similar to that of MCMC flux sampling approaches. Note that FBAwEB is not a method with new types of constraints as in pFBA or ecFBA (Yeo *et al*., 2020), rather it is an extension of FBA and can be incorporated into any constraint-based flux analysis method e.g., pFBA, FVA or ecFBA.

**Figure 3.**
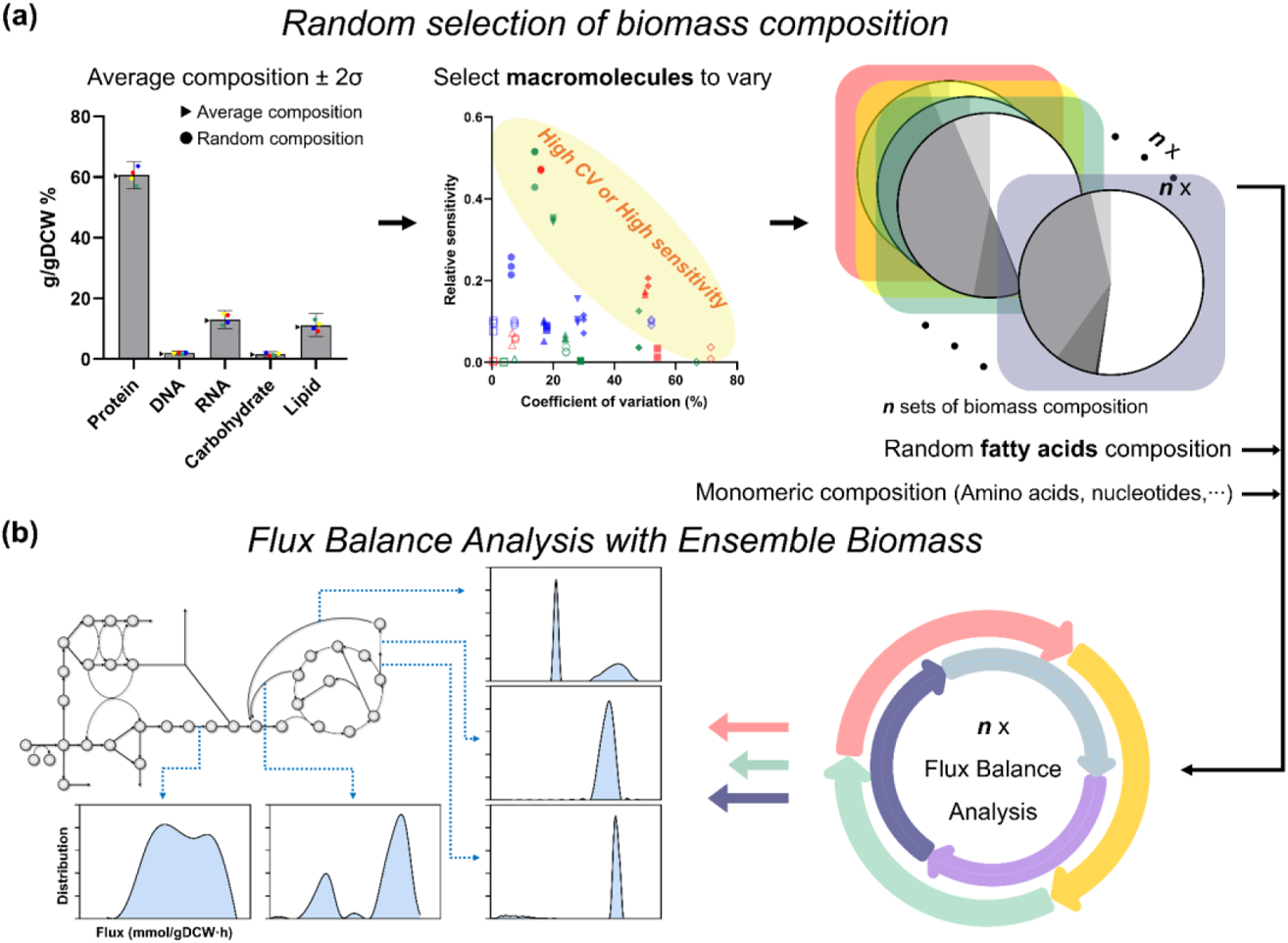
Steps to formulate an ensemble biomass and process data of Flux Balance Analysis with Ensemble Biomass (FBAwEB). First step of FBAwEB is to determine a single set of ‘reference’ macromolecular/monomer compositions, which can be determined based on experimental data or data of phylogenetically-close organisms. This ‘reference’ will be the standard composition before compositional changes are taken. Based on the coefficient of variation (CV) of each macromolecule estimated in this work or known variable ranges, ***n*** sets of macromolecular compositions can be obtained. Monomer compositions of amino acids, deoxyribonucleotides, ribonucleotides, etc., remain constant. With every single macromolecular composition, a single set of fatty acids composition is randomly (but within a certain range) determined and one set of biomass-related equations is formulated. In this way, FBA is iteratively conducted with Ensemble biomass including ***n*** sets of biomass equations. As a result, flux distributions of all reactions can be obtained.

To demonstrate the applicability of newly proposed approach, we implemented pFBA with Ensemble biomass (pFBAwEB) using the GEMs of *E. coli*, *S. cerevisiae* and CHO cells, and compared its performance with pFBA using relevant experimentally measured flux data determined by carbon isotope labelling (Chen *et al*., 2011; Wasylenko and Stephanopoulos, 2015; Templeton *et al*., 2014). Overall, the correlation coefficients of median flux values obtained from the distribution of flux samples from pFBAwEB with corresponding C13MFA fluxes was very similar to that of ones obtained between pFBA and C13MFA (**Figure 4a-c**). However, a closer examination of individual reaction fluxes predicted by pFBAwEB and pFBA indicated substantial differences. Since pFBAwEB predicts a range of values for each reaction flux than a single value as it is in pFBA, it could well represent the uncertainty in some of the fluxes as observed in C13 MFA results, especially in case of *E. coli* and CHO cells (**Figure 4d, Supplementary figure 5**). For example, in *E. coli* aerobic condition, no flux was predicted by pFBA through the reaction catalysing the conversion of 5,10-Methylenetetrahydrofolate (mlthf) to CO_2_ (amino acid metabolism) although it was experimentally determined to carry flux. On the other hand, pFBAwEB predicted the flux values within the measured range. Moreover, the comparison among C13 MFA, pFBAwEB, and pFBA implied two main strengths of FBAwEB. First, compared to pFBA, pFBAwEB presents biologically relevant flux ranges by exploring uncertainty in biomass compositions. It should be noted that this range of solutions obtained using pFBAwEB accounts for natural variation in cellular organisms rather than solely relying on mathematical or statistical techniques. Next, despite using an ensemble biomass, the predicted flux spans generally did not surpass the extent of errors of C13 MFA data and presented distinguishable patterns when compared between conditions. For example, the flux ranges predicted in *E. coli*-aerobic was significantly different from *E. coli*-anaerobic condition (**Figure 4d**).

**Figure 4.**
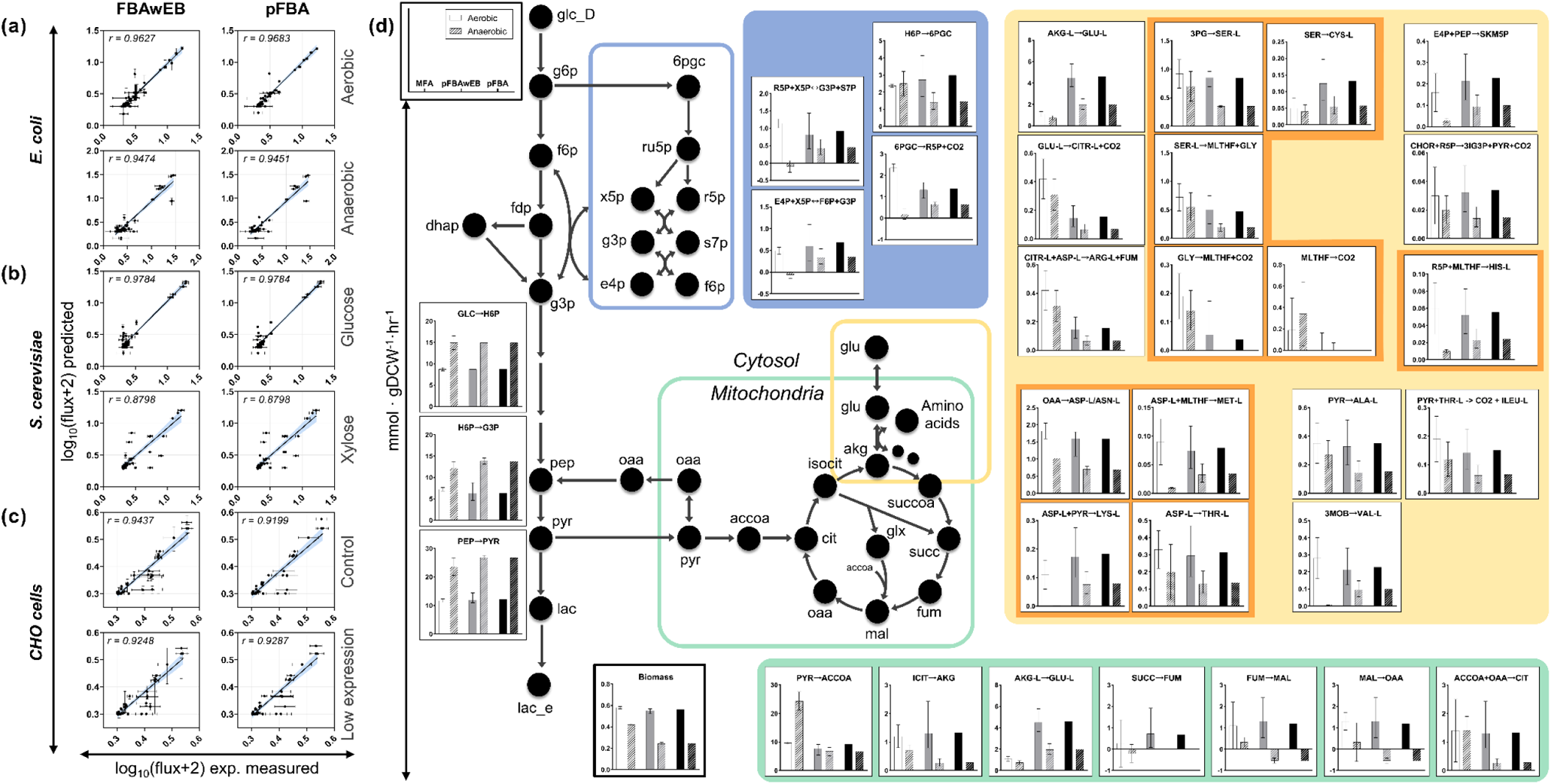
Comparison of flux distributions with and without ensemble biomass (pFBAwEB vs pFBA). **4a-c**. Pearson correlation between the experimentally measured fluxes and the predicted fluxes by FBAwEB and pFBA under two different conditions each (*E. coli* with aerobic/anaerobic conditions; *S. cerevisiae* with glucose/xylose uptake conditions; CHO cells with control/low expression of Bcl-2Δ conditions). **4d**. Detailed comparison of experimentally estimated fluxes (13C MFA, white bar) with the predicted fluxes by pFBAwEB (grey bar) and pFBA (black bar). The central carbon metabolism including PPP (Pentose Phosphate Pathway, blue box) and TCA cycle (green box), and various amino acids metabolism (yellow box) of *E. coli* under aerobic (solid bar) and anaerobic (pattern bar) are represented. The prediction by pFBAwEB can present biologically-relevant flux ranges which cannot be obtained by pFBA by resolving biomass uncertainty.

## 3. Discussion

Understanding the degree of natural variation in macromolecules and monomers is fundamental for mitigating the uncertainties in biomass equations and their influence in intracellular flux predictions using FBA. The current study investigated the natural variations in biomass of three representative organisms and gives a comprehensive overview on how cellular composition vary across different conditions. While most macromolecules varied significantly, composition of monomers, except fatty acids, remain relatively stable across conditions. CV values of carbohydrate, DNA, and RNA were much greater than that of their corresponding monomers while protein and lipid showed less variability than their monomers in most cases. Considering that the mean value and the CV are in an inverse relation, relatively small CV values of protein compared to other macromolecules are reasonable because protein accounts for the largest amount of cell weight (approx. 50% mass fraction). Similarly, very little amount of carbohydrate in *E. coli* (1.6 % of gDCW) would have contributed to substantially high CV. Unlike macromolecules, composition of monomers varied relatively lesser across different conditions, except fatty acids. These observations together clearly indicate the necessity of accurate measurements of various biomass components, particularly the macromolecular composition. In this regard, Beck *et al*. (2018) reviewed multiple analytical techniques and suggested guidelines for accurate quantitative measurement of five major macromolecules for biomass equations. While FBAwEB can be directly applied to an organism with multiple biomass measurements available, we also suggest it can be applied to organisms which doesn’t have multiple biomass measurements across conditions. Biomass compositions measured in a singular condition in such instances can be used as reference and the range of CVs reported in this study shall be adopted to account for the natural variation in biomass when appropriate data is not available. This suggestion stems from the observation that phylogenetically close species showed similar trends in macromolecular distribution.

We also examined how reliable it is to estimate biomass components from omics datasets. We noted there exists difference between experimental and estimated monomer compositions from omics data (**Figure 1d**), which may stem from two main reasons. Firstly, these methods are based on an idealistic assumption regarding gene and protein expression. In particular, ribonucleotide composition estimated from genome sequence assumes that all genes in the genome are transcribed, but in reality, only around 5% of the genome is transcribed into RNA at any given time (Frith *et al*., 2005), and thus using genomic data for ribonucleotide and amino acid composition estimation is inappropriate. Moreover, even with transcriptome data that represents gene expression, this only takes in account the mRNA, rRNA and other small RNA portion of total RNA content, but usually does not cover the more abundant tRNA. Similarly, estimation of amino acid composition from the genome contains an underlying assumption that all proteins are always expressed, which is clearly untrue. In agreement with our results, highly expressed transcriptomic and proteomic data can provide a better estimation. However, we still noted minor variations between the omics-data derived estimates and the experimentally measured compositions which could be due to local variations. Notably, nucleotide, ribonucleotide and amino acid compositions are known to vary locally in individual members of a population. In particular, nucleotide composition varies locally in different areas of a given genome (Bohlin *et al*., 2010), and consequently could result in mRNA transcripts with varying ribonucleotide composition. Similarly, amino acid composition differs across protein functional categories, probably related to consideration of translation rate (Akashi and Gojobori, 2002). Fortunately, even though omics-data estimated monomer composition varies marginally from experimental values, our sensitivity analysis results showed that while ribonucleotide composition does not impact intracellular flux distribution significantly, amino acid composition influences only moderately (**Figure 2**). Therefore, we conclude that using estimated ribonucleotide and amino acid values, preferably from highly expressed transcriptome or proteome data in the biomass equation is a reasonable approach.

Apart from understanding the natural variation of biomass components, understanding the sensitivity of intracellular flux distributions to biomass components is also critical. In this regard our analysis indicated that flux sensitivity was dependent on both organisms’ metabolic network and environment and/or genetic conditions (**Figure 2**). For example, while absolute sensitivities of *E. coli* and *S. cerevisiae* were smaller and mostly insignificant, it was much more pronounced in CHO cells. This discrepancy in the sensitivities between organisms is mainly attributed to the metabolic network structure as described previously (Yuan *et al*., 2016). In addition, absolute sensitivities were divergent between different environmental conditions for same species. Nonetheless, the relative sensitivities indicated that, although the absolute consequence of biomass variations may fluctuate depending on model conditions, its relative influence on flux distribution remains consistent across organisms. Protein was consistently identified as the component with high relative sensitivities among all macromolecular classes and monomers.

A few efforts were recently undertaken to address the uncertainty of biomass composition within or across species before our work (Bernstein *et al*., 2021). In one such attempt, known as BOFdat, a random sampling method was used to obtain a biomass equation that best-fit the known gene essentiality data (Lachance *et al*., 2019). While BOFdat implemented random sampling of biomass equations similar to our approach, it does not account for the variation in macromolecular composition which we have observed to vary the most across conditions. Instead, BOFdat varies individual components of the biomass equation in a binary fashion by including or excluding it in the biomass equation focussing only on the essentiality prediction and ignores the sensitivities of intracellular fluxes. Another work suggested two frameworks for addressing condition-specific variations of biomass compositions upon nutritional changes: 1) an optimal set of trade-off weights assigned to multiple biomass equations is chosen for the maximal growth rate, 2) the biomass composition is estimated by interpolation based on known sets of compositional data measured under different environmental conditions, assuming linearity between compositions and the environmental changes (Schulz *et al*., 2021). Although this approach attempts to account for natural variation in the biomass composition, the key limitation of this approach lies on its theoretical treatment of biomass variations, i.e., linear variation in environments. It also just relies on two different biomass measurements and if these two experiments do not cover the extremities of the biomass variation then the interpolation is applicable only within a subspace of potential range. In this regard, FBAwEB address the above-mentioned shortcomings by relying on experimentally observed natural variation in biomass components from multiple datapoints and treats the variation within the observed form in a stochastic manner in the form of ensembles.

Overall, we conclude that among various components of biomass, all macromolecules and fatty acids vary considerably under different environments. Thus, these need to be accounted for its natural variation carefully while drafting the biomass equation. We also found that the omics data-estimated composition of monomers, particularly amino acids, is within the observed natural variation and could be reliably used. Based on such observations, our new approach, FBAwEB, can account for both the natural variations in biomass compositions and their intracellular flux sensitivities. This will facilitate more reliable and accurate predictions of metabolic states and physiological behaviours in GEM guided analysis.

## 4. Material and Methods

### 4.1 Compilation of macromolecule and monomer compositions

Macromolecular and monomer composition data for three representative species, *E. coli*, *S. cerevisiae*, and CHO cells, was collected through a targeted literature search. Both macromolecular and monomer composition data was collected from various conditions including changes in genetic make-up, i.e., mutants, and environmental conditions including different temperature, dilution rates, oxygen concentrations, media compositions, and growth phases. The full list of biomass composition data and its source from where it was obtained are listed in **Supplementary file 2**.

Phylogenetically close organisms of *E. coli*, *S. cerevisiae*, and CHO cells were determined from the realm of Gram-negative bacteria, yeast species, and murine cell or *Mus musculus*. Macromolecular composition data of these organisms was also collected through the same targeted literature search. As complete macromolecular composition data was unavailable for many species, we only included species with at least single dataset of macromolecular composition available in this analysis. The full list of biomass composition data and its source from where it was obtained are listed in **Supplementary file 1**.

### 4.2 Estimation of natural variation in biomass components

The variability of all macromolecules and monomers were calculated using the coefficient of variation (CV). CV of each component was calculated by dividing the standard deviation of collected and/or processed composition data (mass % or mole %) by the mean value of the corresponding biomass component. The CVs of monomers necessitated an additional step here, where the average of multiple monomer CVs was taken to represent a final CV. On the other hand, the variability of monomers of DNA was simply determined based on multiple GC% data crawled with a query of each species name of three organisms from NCBI genome database (Agarwala *et al*., 2017). Note that the estimation of ribonucleotides composition from omics data was not taken into consideration when we evaluated the CVs.

### 4.3 Estimation of monomer composition from omics-data

We collected multi-omics data of *E. coli*, *S. cerevisiae*, and CHO cells to estimate ribonucleotide and amino acid composition. Genome data were obtained from NCBI RefSeq database (O’Leary *et al*., 2016), transcriptomic datasets were collected from the Sequence Read Archive (Leinonen *et al*., 2011) and NCBI Gene Expression Omnibus (GEO), and proteomic data was downloaded from PaxDB (Wang *et al*., 2012). The source information of each omics data used in this study is listed in **Table 1**. Note that amino acid composition is not estimated from proteome for CHO cells due to lack of whole proteomic data in PaxDB.

**Table 1.**
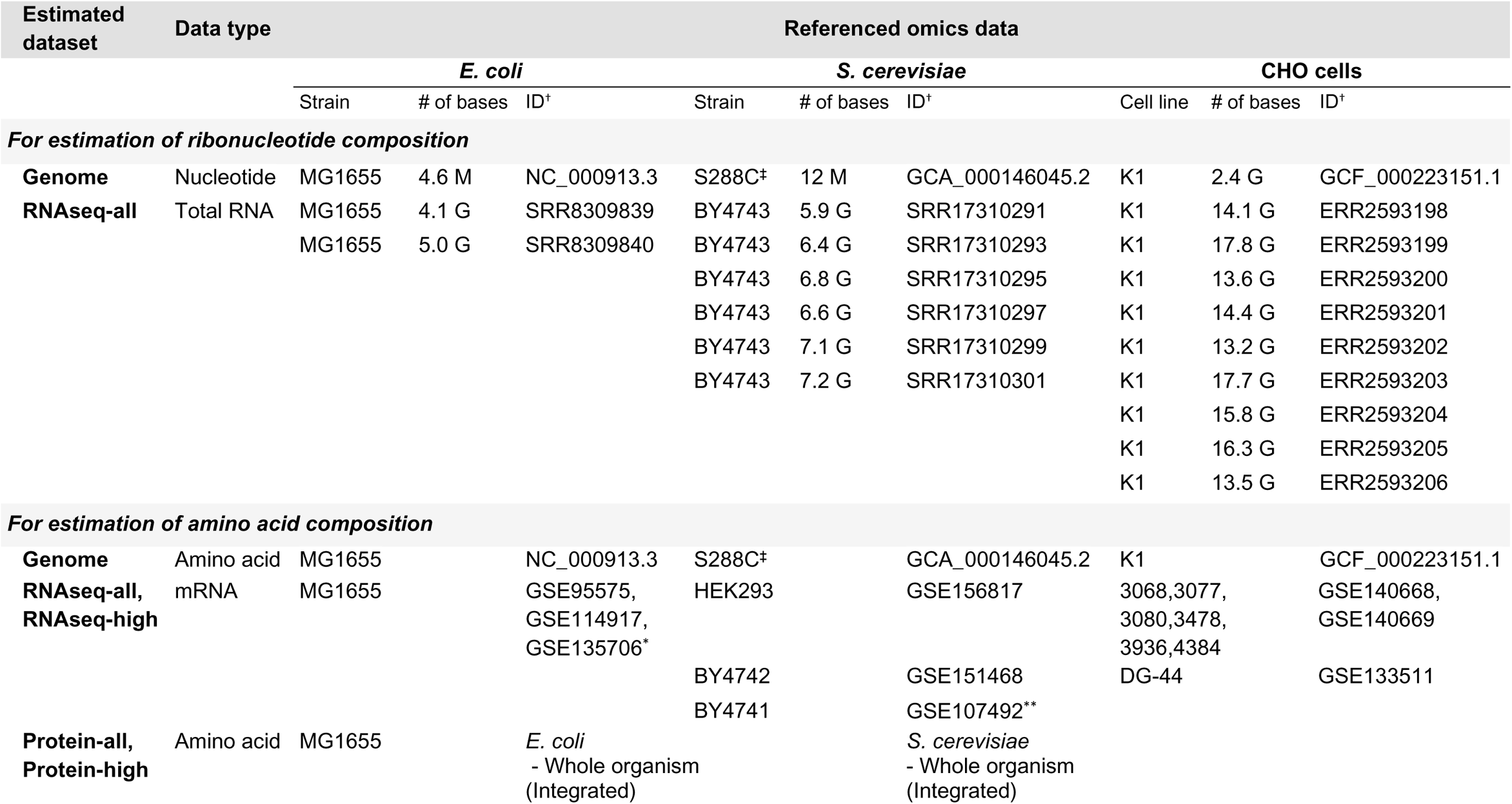
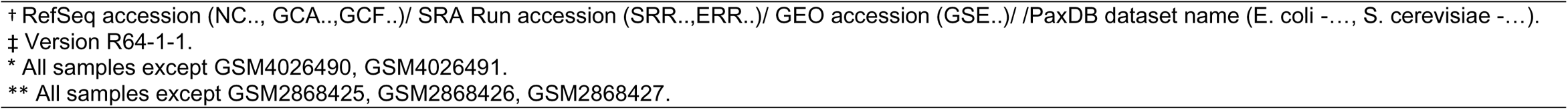
Information of omics data used for estimation of ribonucleotide/amino acid composition.

For the transcriptome and proteome data, gene lists with their expression values were used to extract corresponding coding sequences to estimate the ribonucleotide and amino acid compositions. These gene lists were classified into two categories: all expressed and top 10% highly expressed genes. Note that we excluded some datasets from the GEO series which were obtained from environmental/genetic conditions that resulted in reduction of cellular growth more than 30% when compared to control conditions. In case of genome data, all known genes encoded were considered. These gene sets were labelled according to their respective source: Genome, RNAseq-all, RNAseq-high, Protein-all, and Protein-high. Ribonucleotide composition was estimated from Genome, RNAseq-all and RNAseq-high sequences. While the coding sequences of genes were considered as it is for genome and transcriptome data, a corresponding amino acid sequence was first obtained using the ExPasy translate tool. Then the frequency of monomer 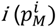 is obtained from the coding sequences as follows:

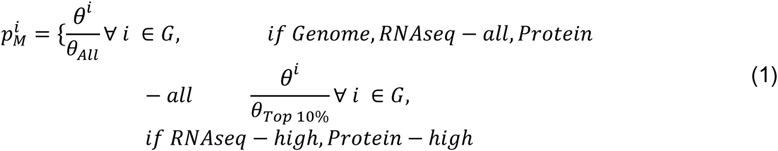

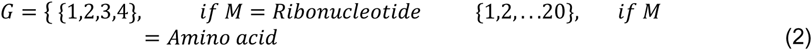

where *θ^i^* stands for the occurrence of monomer *i*, and *θ_All_* is the total number of monomers. Total number of monomers in genome was based on coding sequences of all genes while in transcriptome and proteome data it was based on expressed genes only. *θ_Top 10%_* denotes the total number of monomer units encoded by top 10% expressed genes in transcriptome or proteome data while *θ_All_* considers all genes expressed (count >10).

### 4.4 Flux balance analysis and sensitivity of biomass composition on intracellular flux distribution

The following GEMs were used for each organism: *E. coli* - *i*ML1515 (Monk *et al*., 2017), *S. cerevisiae* - Yeast8.0.0 (Lu *et al*., 2019) and CHO cells - *i*CHO2291 (Yeo *et al*., 2020). pFBA was conducted in the entire work and its formulation is represented as follows:

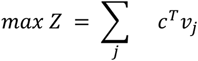

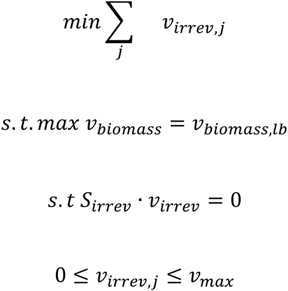

where Z is an objective function, *c^T^* is a weight vector of reactions contributing to the objective function, ν*_j_* is a flux of reaction j, *S_irrev_* is a stoichiometric matrix where all reactions are represented as irreversible, and *ν_biomass,lb_* is the lower bound for the biomass synthesis reaction.

In order to test the sensitivity of flux predictions to biomass composition, we altered the target component in biomass equation by 25% of the average mass fraction (g/gDCW) followed by normalization so that the sum of all components became 1gDCW, while maintaining the original relative amounts of other macromolecules. The monomer compositions were kept at the original values while varying the macromolecular mass fractions, and vice versa. The biomass equation (**B**) can be represented as follows:

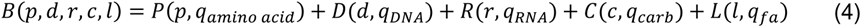

where p, d, r, c, l denotes the macromolecular weight fraction of protein, DNA, RNA, carbohydrate, and lipid, respectively. ***P***, ***D***, ***R***, ***C***, ***L*** are macromolecular synthesis equations which are functions of q monomer composition vectors. A biomass equation with altered macromolecular composition, e.g., protein, will be

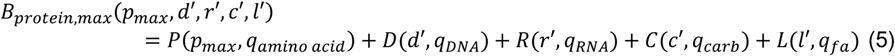

where *p_max_* represents the protein fraction increased by 25%, and d’, r’, c’, l’ denote the corresponding normalized values.

Here, we sought to estimate the sensitivity coefficients of each biomass component using Euclidean distances (ED) between the fluxes obtained without altering the biomass equation, i.e. reference condition, to that of altered biomass equations. Briefly, a flux vector ν of any biomass equation is obtained by performing pFBA. Subsequently, the flux vectors (ν) were normalized to the length of each vector. Finally, ED is calculated as the root mean square of the difference between the two vectors. Mathematically,

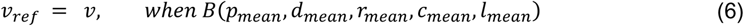

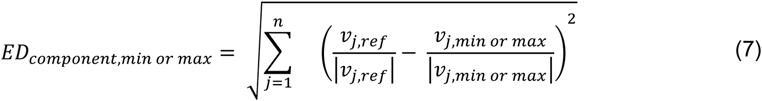

where ***component*** denotes macromolecule class (e.g., protein or DNA) and all members of monomers (e.g., alanine or glycine of protein, and dATP or dCTP of DNA), ***n*** is the number of reactions in a GEM.

The average of these EDs was used as the definition of the (absolute) sensitivity,

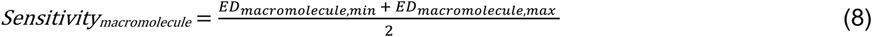

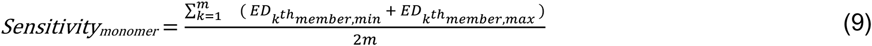

where ***macromolecule*** is protein, DNA, RNA, carbohydrate, and lipid, ***monomer*** represents monomer groups such as amino acids, deoxyribonucleotides, ribonucleotides, and fatty acids, and ***m*** is the number of members consisting each monomer group. For example, the *Sensitivity_protein_* is obtained by taking an average of *ED_protein,min_* and *ED_protein,max_*, and *Sensitivity_amino acids_* is an average value of *ED_alanine,min_*, *ED_alanine,max,_ ED_glycine,min_*, *ED_glycine,max_*, and so on. A relative sensitivity was obtained by normalizing an absolute sensitivity by the sum of the absolute sensitivities of all macromolecules and monomers for every model condition.

### 4.5 Principal component analysis

Using maximum and minimum composition of each macromolecule and monomer component as stoichiometric coefficients of biomass equations, we implemented pFBA to obtain flux distributions for principal component analysis. We excluded several reactions which had zero fluxes on each sample (biomass equation) and normalized them with the resultant flux distribution obtained from the reference condition which takes the mean composition of each macromolecule and monomer component as stoichiometric coefficients. Each variance ratio and principal component scores were calculated based on the processed data using scikit-learn (Pedregosa *et al*., 2011) and Matplotlib package (Hunter, 2007) by Python 3.8.0.

### 4.6 Statistical analysis

Kruskal-Waliis H test (Kruskal and Wallis, 1952), which is a rank-based non-parametric method, was used to compare significant differences of macromolecular compositions between *E. coli*, *S. cerevisiae*, CHO cells, and their relatives (**Figure 1c**). Sensitivities of biomass components were compared using one-way ANOVA corrected for a multiple comparisons test with the two-stage step-up method of Benjamini, Krieger and Yekutieli (Benjamini *et al*., 2006). This method is one of the false discovery rate (FDR) approaches for multiple comparisons. The FDR-adjusted p values (q values) were obtained with the desired false discovery rate of 0.05. Comparison between the distance distribution of EXP_n_-EXP_m_ and (RNA-seq or Protein high)_n_-EXP_n_ was conducted by Mann-Whitney U test (Mann and Whitney, 1947). All statistical tests in this work were performed by GraphPad Prism version 9.3.1 for Windows, GraphPad Software, La Jolla California USA, www.graphpad.com.

### 4.7 Flux Balance Analysis with Ensemble Biomass (FBAwEB)

For all three models, the synthesis equation of each macromolecule was re-formulated based on the original biomass equations in the model of Yeast8.0.0., which has individual macromolecular synthesis equations. For example, as the biomass equation of *i*ML1515 is not subdivided into model macromolecular metabolites such as ‘protein[c]’, ‘DNA[c]’, ‘RNA[c]’, ‘carbohydrate[c]’, ‘lipid[c]’, we added these macromolecular metabolites and reallocated monomers into the corresponding macromolecules in all models. Based on summation of all model stoichiometric coefficients (mmol/gDCW) of compounds in lipid synthesis reaction and total lipid weight (g/gDCW) used for Yeast8 model (Lahtvee *et al*., 2017), we estimated the molecular weight (MW) of total lipid to be 3100 g/mol and assume the MW of lipid is constant under varying conditions. We also added a fatty acid synthesis reaction based on the abovementioned data to specify fatty acids composition. For every 5000 iterations, we randomly re-distributed macromolecular and fatty acids composition within a range within ± 2×σ (standard deviation determined based on the collected experimental data) from a reference composition. The reference compositions are classified into two categories: Macromolecular and monomer composition. All macromolecular reference compositions were based on the average of multiple collected experimental data. On the other hand, several reference compositions for lipid compounds (for *S. cerevisiae* model) and fatty acids (for all three models) were obtained from a single composition of the original model due to non-uniform data structures in various data sources. Moreover, since the difference of growth associated with maintenance (GAM) exhibits a distinct effect on model-predicted growth rate of different organisms (Yuan *et al*., 2016), we fixed the maintenance cost for each GEM in order to understand the consequences of compositional variation excluding the effect of maintenance energy cost.

We collected the experimentally determined intracellular flux data for *E. coli*, *S. cerevisiae*, and CHO cells (Holm *et al*., 2010; Blank *et al*., 2005; Templeton *et al*., 2014) to compare it with the intracellular fluxes predicted by pFBA and pFBAwEB. Note that pFBA and FBAwEB simulations were performed reflecting the experimental conditions whose isotope labelled flux data were available. In this study, pFBA and pFBAwEB was implemented using COBRA Toolbox 3.0 (Heirendt *et al*., 2019) within the MATLAB (The MathWorks, version R2020a) environment and Gurobi solver (http://www.gurobi.com, version 9.1.1) to solve the underlying optimization problem.

## Availability of data and materials

GEMs, MATLAB and python scripts and corresponding datasets generated during the study are available at the GitHub repository: https://github.com/skku-pdse/FBAwEB.git

## Author contributions

Y.-M. Choi: Methodology, Formal analysis, Software, Visualization, Writing - Original Draft; D.-H. Choi: Software, Writing - Original Draft; Y. Q. Lee: Data Curation; L. Koduru: Methodology, Writing - Review & Editing; N. E. Lewis: Conceptualization, Writing - Review & Editing; M. Lakshmanan: Conceptualization, Methodology, Writing - Review & Editing, Supervision; D.-Y. Lee: Conceptualization, Writing - Review & Editing, Supervision.

## Acknowledgements

We would like to thank Su Kyung Kim, Minouk Lee, and Dongseok Kim for assistance in collecting relevant omics data. D.Y.L. acknowledges funding support from the A*STAR Research Attachment Programme (ARAP), Singapore, the Korea Health Technology R&D Project through the Korea Health Industry Development Institute (KHIDI) funded by the Ministry of Health & Welfare, Republic of Korea [grant number HI19C1348], and the Korea Institute of Planning and Evaluation for Technology in Food, Agriculture, Forestry and Fisheries (iPET) through MAFRA programme [grant number 32136-05-1-HD050]. N.E.L. acknowledges funding support from National Institute of General Medical Sciences, United States [grant number R35 GM119850], and the Novo Nordisk Foundation [grant number NNF20SA0066621]. M. L. acknowledges funding support from A*STAR, Singapore through IAF-PP (HBMS) programme [grant number: H20H8a0003].

