## Supplementary Figure for "Mitigating biomass composition uncertainties in flux balance analysis using ensemble representations"

**Supplementary figures**

**Supplementary figure 1**. Growth rate changes at maximum and minimal composition (determined based on collected data) of each biomass component under various conditions.

**Supplementary Figure 2**. PCA results of relative flux changes of all metabolic reactions. between flux distributions of reference composition and maximum/minimum composition of macromolecules or monomers. For *E. coli*, flux distributions under aerobic and anaerobic conditions are implemented. Likewise, 3 different conditions of *S. cerevisiae* under distinct carbon sources (glucose, xylose and ethanol) are considered for flux distribution. Additionally, we constrained CHO GEM with control, Bcl-2Δ low expression (le), and high expression (he) based on the 13C MFA research (Holm *et al.*, 2010; Blank *et al.*, 2005; Templeton *et al.*, 2014).

**Supplementary Figure 3**. Relative sensitivities of biomass components on flux prediction and p-values between biomass component groups. The p-values are the result of a multiple comparison test with the two-stage step-up method of Benjamini, Krieger and Yekutieli (the desired false discovery rate of 0.05). Only q-values (FDR-adjusted p values) less than 0.05 are represented.

**Supplementary Figure 4**. Detailed comparison of experimentally estimated fluxes (13C MFA, white bar) with the predicted fluxes by pFBAwEB (grey bar) and pFBA (black bar). The conditions of glucose uptake (aerobic) and xylose uptake (aerobic) were used for the S. cerevisiae model.

**Supplementary Figure 5**. Detailed comparison of experimentally estimated fluxes (13C MFA, white bar) with the predicted fluxes by pFBAwEB (grey bar) and pFBA (black bar). The conditions of control state and low expression of Bcl-2Δ condition were used for the CHO cells model.

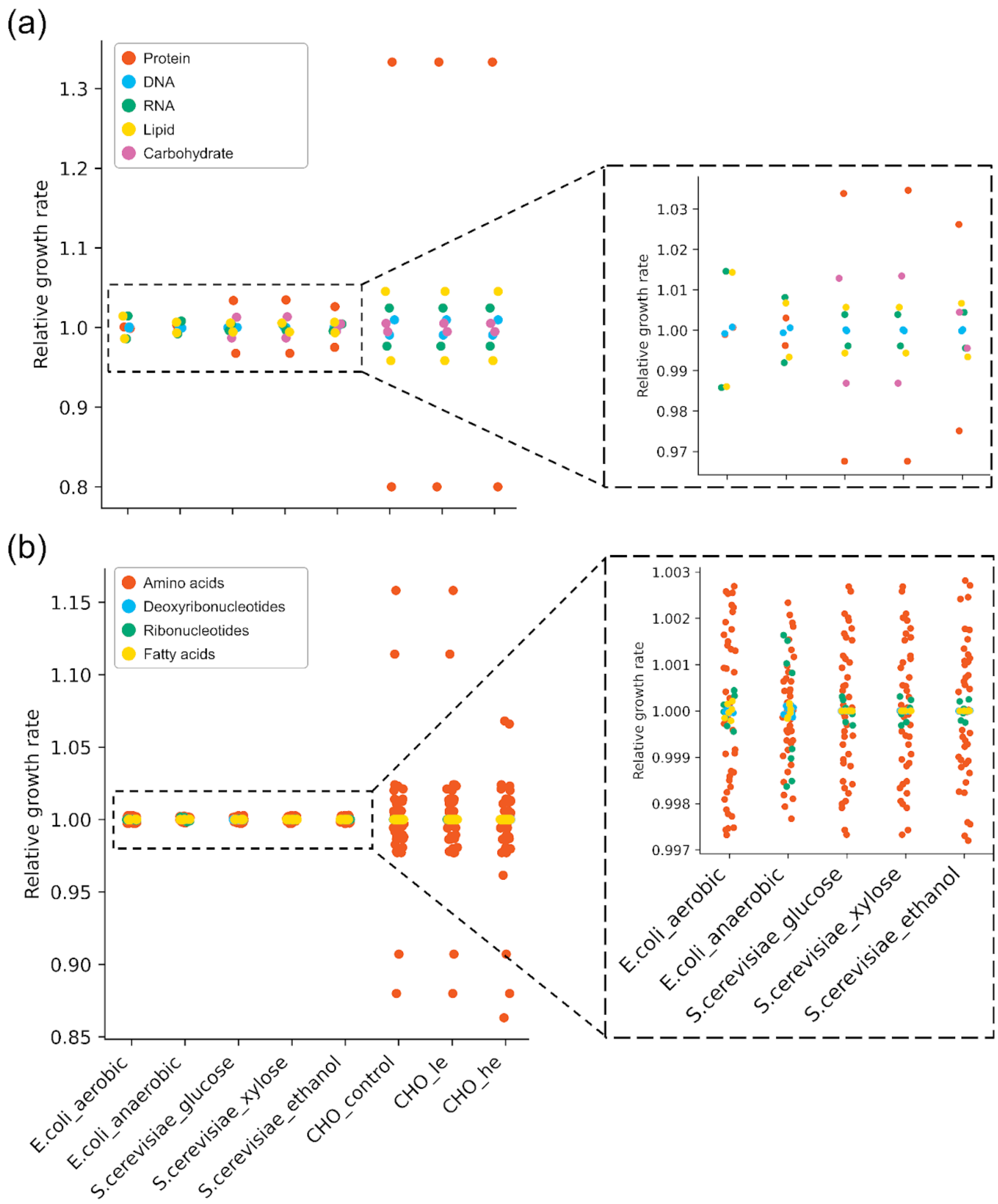


**Supplementary figure 1**. Growth rate changes at maximum and minimal composition (determined based on collected data) of each biomass component under various conditions.

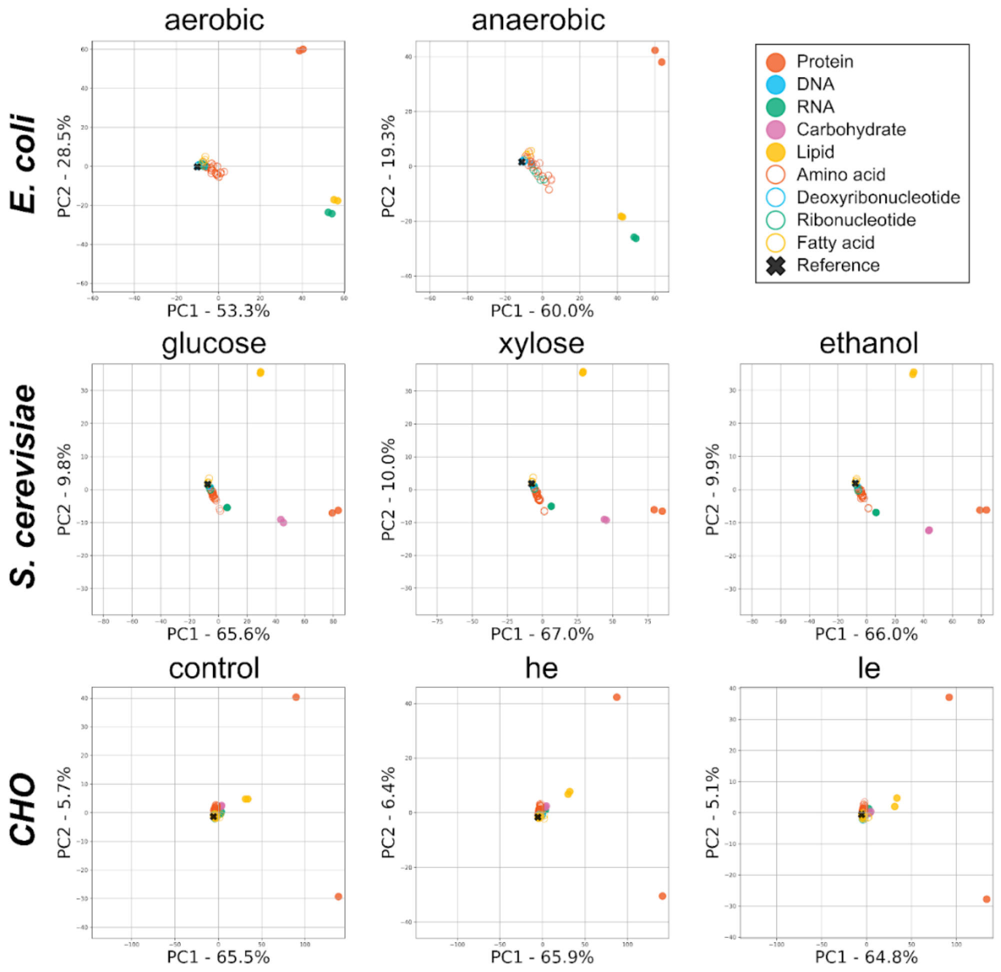


**Supplementary Figure 2**. PCA results of relative flux changes of all metabolic reactions. between flux distributions of reference composition and maximum/minimum composition of macromolecules or monomers. For *E. coli*, flux distributions under aerobic and anaerobic conditions are implemented. Likewise, 3 different conditions of *S. cerevisiae* under distinct carbon sources (glucose, xylose and ethanol) are considered for flux distribution. Additionally, we constrained CHO GEM with control, Bcl-2Δ low expression (le), and high expression (he) based on the 13C MFA research (Holm *et al.*, 2010; Blank *et al.*, 2005; Templeton *et al.*, 2014).

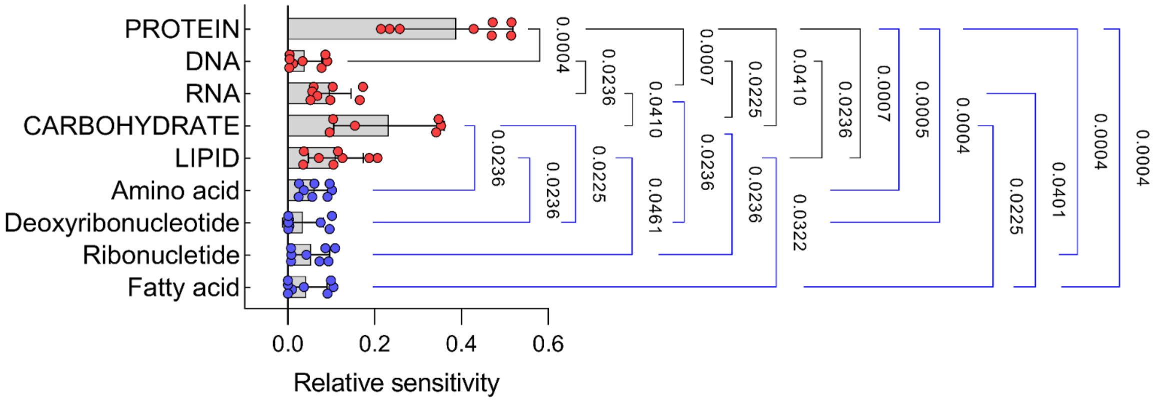


**Supplementary Figure 3**. Relative sensitivities of biomass components on flux prediction and p-values between biomass component groups. The p-values are the result of a multiple comparison test with the two-stage step-up method of Benjamini, Krieger and Yekutieli (the desired false discovery rate of 0.05). Only q-values (FDR-adjusted p values) less than 0.05 are represented.

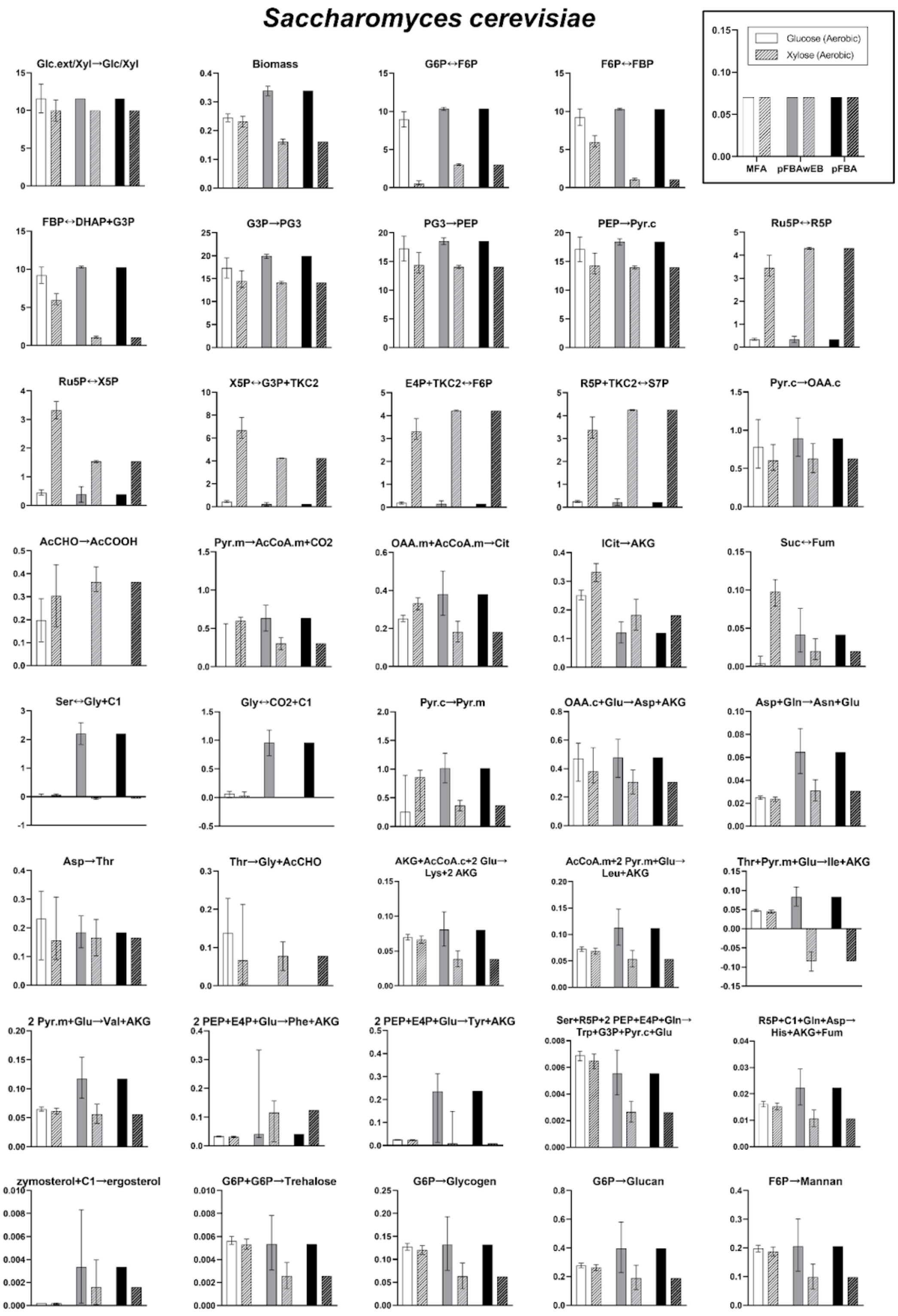


**Supplementary Figure 4**. Detailed comparison of experimentally estimated fluxes (13C MFA, white bar) with the predicted fluxes by pFBAwEB (grey bar) and pFBA (black bar). The conditions of glucose uptake (aerobic) and xylose uptake (aerobic) were used for the S. cerevisiae model.

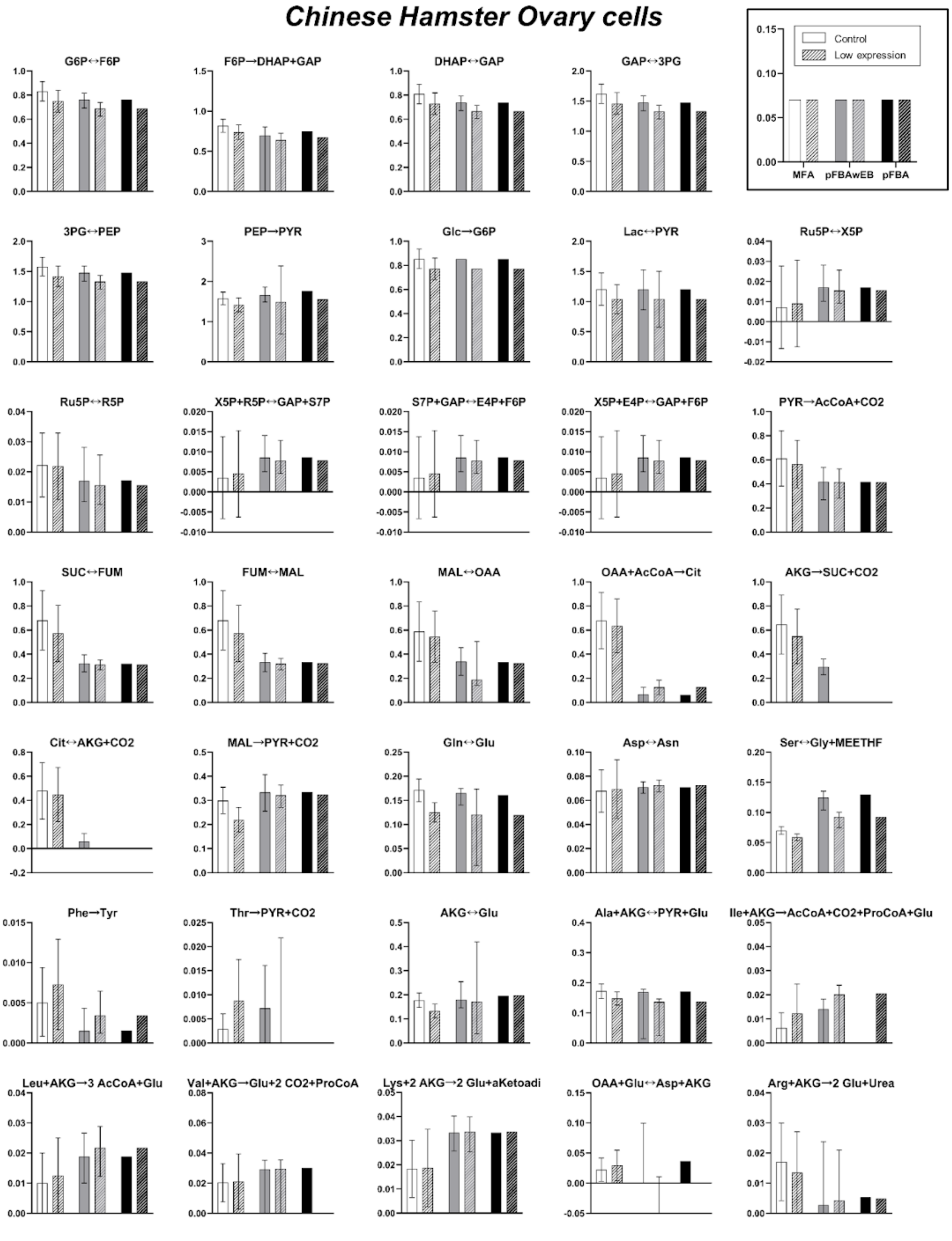


**Supplementary Figure 5**. Detailed comparison of experimentally estimated fluxes (13C MFA, white bar) with the predicted fluxes by pFBAwEB (grey bar) and pFBA (black bar). The conditions of control state and low expression of Bcl-2Δ condition were used for the CHO cells model.
